# Learning can be detrimental for a parasitic wasp

**DOI:** 10.1101/2020.08.17.253641

**Authors:** Valeria Bertoldi, Gabriele Rondoni, Ezio Peri, Eric Conti, Jacques Brodeur

**Affiliations:** Dipartimento di Scienze Agrarie, Alimentari e Ambientali, Università degli Studi di Perugia, Perugia, Italy; Dipartimento di Scienze Agrarie e Forestali, Università degli Studi di Palermo, Palermo, Italy; Département des Sciences Biologiques, Institut de Recherche en Biologie Végétale, Université de Montréal, Montréal, Québec, Canada

## Abstract

Animals have evolved the capacity to learn, and the conventional view is that learning allows individuals to improve foraging decisions. We describe a first case of maladaptive learning where a parasitoid learns to associate chemical cues from an unsuitable host, thereby re-enforcing a reproductive cul-de-sac (evolutionary trap). *Telenomus podisi* parasitizes eggs of the exotic stink bug *Halyomorpha halys* at the same rate as eggs of its coevolved host, *Podisus maculiventris*, but the parasitoid cannot complete its development in the exotic species. We hypothesized that *T. podisi* learns to exploit cues from this non-coevolved species, thereby increasing unsuccessful parasitism rates. We conducted bioassays to compare the responses of naïve *vs*. experienced parasitoids on chemical footprints left by one of the two host species. Both naïve and experienced females showed a higher response to footprints of *P. maculiventris* than of *H. halys*. Furthermore, parasitoids that gained an experience on *H. halys* significantly increased their residence time within the arena and the frequency of re-encounter with the area contaminated by chemical cues. Maladaptive learning in the *T. podisi* - *H. halys* association is expected to further decrease parasitoid reproductive success and have consequences for population dynamics of sympatric native and exotic host species.

## Introduction

Animal decision-making, which is involved in processes such as resource and habitat selection, mate choice and progeny allocation, relies on innate behaviour (instinct), stochastic processes, physiological feedbacks (e.g., hormonal signalling) and learning (reviewed in [1]). Insect parasitoids have been used as model systems to explore both proximate and ultimate perspectives of optimal foraging. In order to cope with spatial and temporal variability in resources, parasitic wasps have evolved the capacity to associate host-related chemical cues to host availability and suitability [2–5]. They further consolidate and improve this capacity through learning processes [6–9], resulting in increased reproductive success [3, 9, 11, 12]. The probability of including a new stimulus in the behavioural repertoire of a parasitoid female depends on its reliability in host location [2], with oviposition having been shown to consolidate a change in foraging behaviour and host acceptance [3, 8, 13]. For example, scelionid parasitoids use host chemical cues on the egg surface or those deposited on plant surfaces by gravid females (footprints) to locate hosts in the habitat (reviewed by [14]). Following detection of host chemical cues, experienced scelionid females show stronger arrestment response (increased residence time, slower walking and increased turning tendency) than naïve females, a behaviour that is re-enforced by successful oviposition [15,16].

Biological invasions generate novel interactions which can have negative consequences on populations of native species [17–20]. This occurs, for example, when an invasive exotic species becomes accepted as a host by native parasitoids but is unsuitable for offspring development. The introduced species then acts as an egg sink [21] for indigenous parasitoids, and negatively impact their reproductive success. We hypothesized that such an evolutionary trap [17, 22, 23] could be exacerbated if foraging parasitoid females learn to exploit cues from a novel but unsuitable host. To our knowledge, there are no examples where associative learning actually results in costs to the foraging success of an animal.

To test this hypothesis we used *Telenomus podisi* Ashmead (Hymenoptera: Scelionidae), a common egg parasitoid of several North American stink bugs (Hemiptera: Pentatomidae) [24, 25]. Abram et al. [26] reported that *T. podisi* females accept and parasitize eggs of the brown marmorated stink bug, *Halyomorpha halys* Stål (Pentatomidae), a recently established pest from eastern Asia [27], at a similar rate to that of its coevolved host, the predator *Podisus maculiventris* (Say) (Pentatomidae). But the parasitoid progeny rarely develop successfully in *H. halys* [26, 28–33], except for a nonconforming *T. podisi* population in California [34]. *Halyomorpha halys* is progressively spreading in invaded areas, thereby increasing the probability of native *T. podisi* encountering this novel but unsuitable host. We examined the learning capacity of *T. podisi* towards its coevolved host, *P. maculiventris,* and non-coevolved host, *H. halys,* and discussed consequences on interacting species.

## Material and methods

### Hosts and parasitoid

A colony of *H. halys* reared continuously on raw pumpkin seeds, carrots, green beans, grapes and potted soybean plants, was established from adults collected in Ontario (Canada) in 2012. The *P. maculiventris* colony was initially established using adults collected in Ontario in 2011 and 2012 and supplemented with bugs from Anatis Bioprotection (Canada). They were fed with live mealworm, *Tenebrio molitor* L. (Coleoptera: Tenebrionidae) larvae, fresh green beans and bean plants. Nymphs were kept in plastic cylinders and fed with mealworm and green beans. For both stink bug species, freshly laid eggs (< 24 h old) were used for the experiments.

The *T. podisi* colony was established with individuals collected in 2011 and 2012 in Ontario. Adult parasitoids were provided with a 1:1 (vol/vol) honey:water solution rubbed on a small piece of ParaFilm^®^. Each week, 1-2 days-old egg masses of *P. maculiventris* (stuck on filter paper using Pritt® stick glue) were exposed to *T. podisi* females for 24 h to maintain the colony. After emergence, male and female parasitoids were kept together in glass tubes for mating. Naïve (i.e. without oviposition experience), 3-8 days-old females were randomly assigned to the different experimental treatments.

All insects were reared in a growth chamber (Conviron E15) at 24±1 °C, 50±5 percent relative humidity, under a 16L:8D photoperiod.

### Treatments

To determine if *T. podisi* females exhibit learning behaviour the following four treatments were tested: (i) naïve parasitoid females foraging on *H. halys* traces; (ii) females with experience on *H. halys* traces and eggs, then tested on *H. halys* traces; (iii) naïve females foraging on *P. maculiventris* traces; and (iv) females with experience on *P. maculiventris* traces and eggs, then tested on *P. maculiventris* traces. The experiments were conducted from 10:00 to 14:00 at 24±1 °C, 50±5 percent relative humidity, under a 16L:8D photoperiod. Between 39 to 43 replicates were conducted for each treatment.

### Oviposition experience

Experienced parasitoid females were obtained in the following manner. A female of either *H. halys* or *P. maculiventris* in their pre-ovipositional phase (with a physogastric abdomen) was introduced in an experimental arena consisting of a Petri dish (5 cm diam., 1 cm height) placed upside-down on a filter sheath. The Petri dish cover had tissue mesh (0.01 cm holes) to prevent saturation of the atmosphere with volatiles released by the stink bug female. Females walked for 30 minutes on the filter paper to allow contamination with chemical traces. Filter papers soiled by bug’s faeces were discarded. Once the stink bug female was removed, a small egg mass (5-6 eggs) of the same species was placed in the middle of the contaminated area. A naïve (i.e. without foraging and oviposition experience) *T. podisi* female was then introduced in the arena. Once she had oviposited the female was removed, isolated in a 1.5 ml tube for 1 h before being tested as an experienced parasitoid. Females that did not oviposit within 1 h were discarded.

### Parasitoid female response to chemical cues

Following Conti et al. [35], bioassays were conducted on a large filter paper arena (20 × 20 cm) where parasitoid females could move freely on the surface. A 5 cm diam area at the centre was exposed to a female of *H. halys* or *P. maculiventris* in pre-ovipositional phase, as described above. For the assays, a *T. podisi* female was released in the middle of the contaminated area and her behaviour recorded with a HDD video camera (Sony HDR-XR 500) placed 40 cm above the arena. The assay stopped when the wasp left the arena or after 10 min. Individuals that flew away within 10 s after the release were excluded from the analysis (n = 6). Each arena was used to test 5 females.

We recorded the time spent by the female in the arena (total residence time) and the number of times the parasitoids went back to the contaminated area once left (number of re-encounters), indicative of a tortuous path associated with the searching behaviour of a parasitoid female [14, 36, 37].

### Statistical analyses

Generalized linear models (GLMs with Gaussian distribution for time data or Poisson distribution for count data) were fitted to test the effects of host herbivore species (*H. halys* or *P. maculiventris*) and parasitoid previous experience (experienced or naïve) on total parasitoid residence time and number of re-encounters with the contaminated area. Residence time data were subjected to Box-Cox transformation for normalization before the analyses. Significance of the model terms was evaluated by means of F test or Likelihood Ratio Test. Significance of the different variable levels was assessed using the Tukey HSD multiple comparisons procedure, adopting a significance level α = 0.05. Analyses were conducted under R statistical environment [38].

## Results

Total residence time of *T. podisi* differed between treatments (Gaussian GLM: F_(3, 157)_ = 24.96, P < 0.0001). For the *T. podisi - P. maculiventris* association, parasitoid females that had previously experienced the chemical traces left by their native host and had been rewarded with an oviposition did not have a greater residence time in the experimental arena than naïve females. In contrast, females with a rewarded experience on the exotic *H. halys* stayed significantly longer in the arena than naïve wasps when tested on chemical footprints of *H. halys* (Fig. 1A). However, both naïve and experienced parasitoids displayed higher residence time on chemical traces of their coevolved host *P. maculiventris* than those of the exotic *H. halys* (Fig 1A).

**Fig 1.**
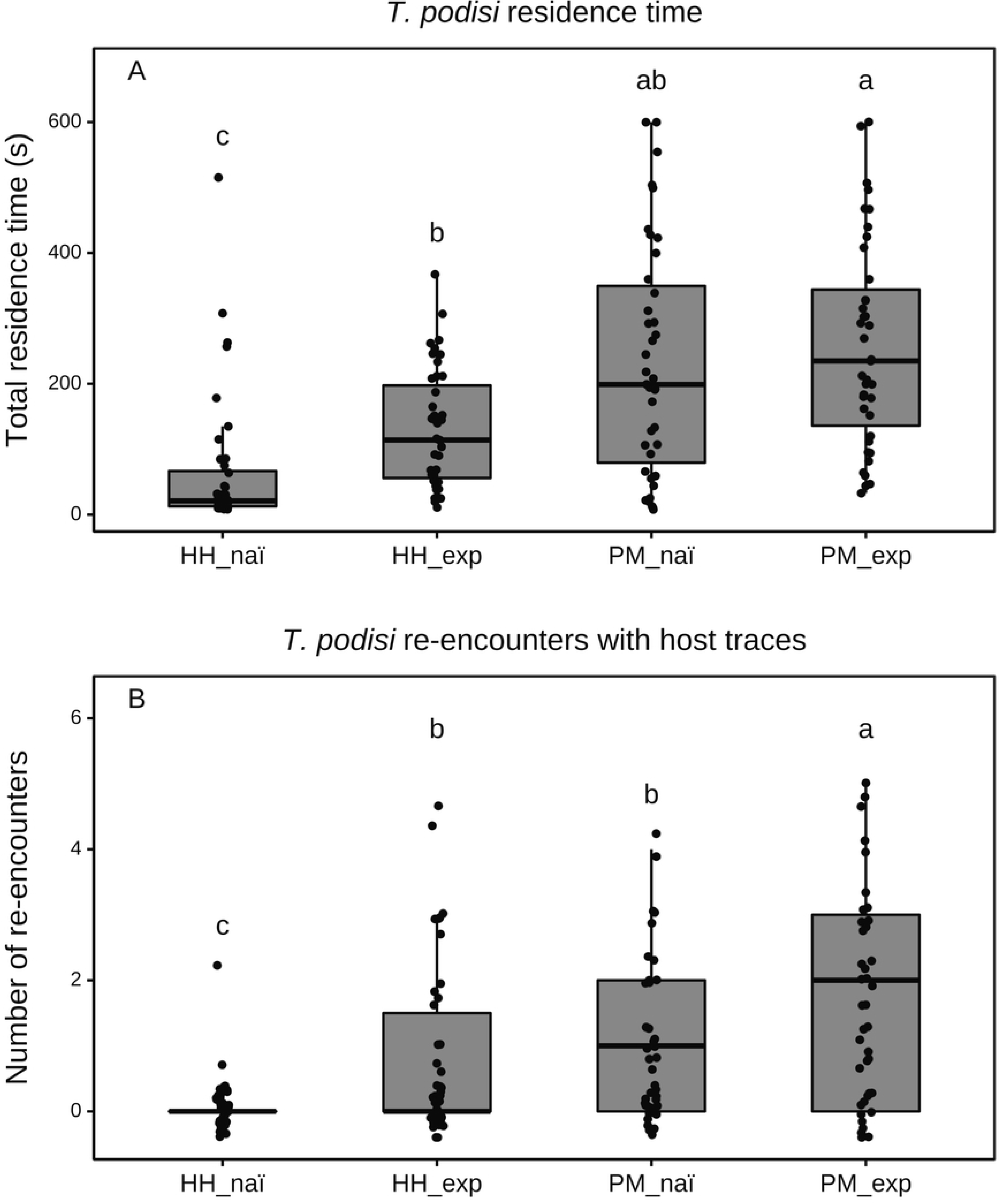
(A) Residence time and (B) number of re-encounters with the host-contaminated arena of naive and experienced wasps. The four treatments were: *T. podisi* naïve females tested on *H. halys* traces (HH_naï; N= 40); *T. podisi* females experienced on *H. halys* traces and eggs, then tested on *H. halys* traces (HH_exp; N= 43); naïve *T. podisi* females tested on *P. maculiventris* traces (PM_naï; N= 39); *T. podisi* females experienced on *P. maculiventris* traces and eggs, then tested on *P. maculiventris* traces (PM_exp; N= 39). Different letters above bars indicate significant differences between treatments (*p* < 0.05; GLM followed by Tukey HSD multiple comparison procedure).

The frequency of re-encounter with the area contaminated by chemical cues differed with parasitoid experience and host species (Poisson GLM: Deviance difference = 74.34, df = 3, P < 0.0001). Experienced *T. podisi* females re-entered the contaminated area more frequently than naïve females (Fig. 1B). Naïve females tested on *H. halys* footprints rarely returned to the contaminated area. However, following a rewarded experience on *H. halys*, the number of re-encounters with this species was similar to the number of re-encounters of naïve females tested on *P. maculiventris* (Fig. 1B). Both naïve and experienced *T. podisi* females returned more frequently to a patch contaminated by *P. maculiventris* than *H. halys* (Fig 1B).

## Discussion

Our findings indicate that *T. podisi* females exhibit increased foraging behaviour following experience with chemical traces and oviposition in either its coevolved and novel host, probably due to associative learning [10, 39]. This capacity permits a female parasitoid to capture and retain information about host availability in the habitat and to adjust her foraging behaviour accordingly. However, the value of learning and its consequences on the reproductive success of *T. podisi* are opposite when exploiting a suitable (*P. maculiventris*) *vs.* unsuitable (*H. halys*) host. Exploiting *H. halys* eggs is maladaptive for *T. podisi* because it incurs significant costs to foraging females and leads to a cul-de-sac for their progeny [26]. *Halyomorpha halys* not only represents an egg sink for *T. podisi*, the most common parasitoid sampled from *H. halys* eggs in Canada [32], but also a ‘time sink’ (sensu [26]) since females increase time foraging in areas contaminated by the unsuitable host. The waste of time is further amplified when females protect the egg mass (patch guarding) from competitors and predators during several hours following oviposition [40]. To our knowledge, this is a first case documenting maladaptive learning in animals.

Both naïve and experienced *T. podisi* females showed a higher response to chemical traces of *P. maculiventris* than of *H. halys*. This pattern was expected because derived stimuli from coevolved hosts are likely to evoke strong innate responses by naïve individuals; indeed the innate response can be stronger than the learned response in the location and acceptance of highly suitable host [41]. However, we cannot exclude an effect of the rearing host (*P. maculiventris*), as shown in other parasitoid species [42–44]. For instance, similar residence time between naïve *vs.* experienced wasps when exposed to *P. maculiventris* chemical traces could be partly explained by *T. podisi* females having already gained experience when developing in and emerging from *P. maculiventris* eggs.

Associative learning of cues from a novel but unsuitable host would also exacerbate the negative effects of the evolutionary trap on *T. podisi* populations. It may contribute to modify the community structure in areas invaded by *H. halys* through direct and indirect ecological effects [26, 45]. Recent exposure to *H. halys* may lead to an increase in *T. podisi*’s rate of parasitism on the invasive host (host switching), and a consequent decrease in parasitism of indigenous pentatomid species (indirect apparent competition). Additional research is required to determine the extent to which detrimental learning would affect the population dynamics of native stink bugs and parasitoids. We only tested females 1 h after their experience with host chemical cues and it would be important to determine how long they exhibit such learned behaviour under natural conditions, considering ecological factors such as the relative densities of native and exotic stink bugs and the persistence of the chemical cues in the footprints.

This original case of maladaptive learning arises from a situation where a native parasitoid encounters a new potential host species as a result of a biological invasion. On one hand, there is no operational ecological filter that stops host location and acceptance and, on the other hand, there is a strong physiological filter that prevents parasitoid development [31]. Accordingly, a parasitoid could escape such an evolutionary trap by evolving (i) behavioural capacities to prevent acceptance of an unsuitable resource or (ii) physiological capacities to successfully reproduce in the novel host species [26, 46].

## Acknowledgments

We thank Paul K. Abram and Jeremy N. McNeil for helpful comments and Josée Doyon for technical assistance.

## References

1. Papaj DR, Lewis AC. Insect learning: Ecology and evolutionary perspectives. London, UK: Chapman & Hall; 1993.

2. Vet LEM, Dicke M. Ecology of infochemical use by natural enemies in a tritrophic context. Annu. Rev. Entomol. 1992; 37: 141–172. doi : 10.1146/annurev.en.37.010192.001041

3. Turlings TC, Wäckers FL, Vet LEM, Lewis WJ, Tumlinson JH. Learning of host-finding cues by hymenopterous parasitoids. In: Papaj DR, Lewis, AC, editors. Insect learning. London, UK : Chapman; 1993. pp. 51–78. doi : 10.1007/978-1-4615-2814-2

4. Godfray HCJ. Parasitoids: behavioral and evolutionary ecology. New Jersey, USA : Princeton University Press; 1994.

5. Fatouros NE, Dicke M, Mumm R, Meiners T, Hilker M. Foraging behavior of egg parasitoids exploiting chemical information. Behav Ecol. 2008; 19: 677–689. doi : 10.1093/beheco/arn011

6. Arthur AP. Associative learning by *Nemeritis canescens* (Hymenoptera: Ichneumonidae). Can Entomol. 1971; 103: 1137–1141. doi : 10.4039/ent1031137-8

7. Lewis W, Tumlinson JH. Host detection by chemically mediated associative learning in a parasitic wasp. Nature. 1988; 331: 257. doi : 10.1038/331257a0

8. Vet LEM, Groenewold AW. Semiochemicals and learning in parasitoids. J Chem Ecol. 1990; 16: 3119–3135. doi : 10.1007/bf00979615

9. Dukas R. Evolutionary biology of insect learning. Annu Rev Entomol. 2008; 53: 145–160. doi : 10.1146/annurev.ento.53.103106.093343

10. Papaj DR, Prokopy RJ. Ecological and evolutionary aspects of learning in phytophagous insects. Annu Rev Entomol. 1989; 34: 315–350. doi : 10.1146/annurev.en.34.010189.001531

11. Dukas R, Duan JJ. Potential fitness consequences of associative learning in a parasitoid wasp. Behav Ecol. 2000; 11: 536–543. doi : 10.1093/beheco/11.5.536

12. Nieberding CM, Van Dyck H, Chiitka L. Adaptive learning in non-social insects: from theory to field work, and back. Curr Opin Insect Sci. 2018; 27: 75–81. doi : 10.1016/j.cois.2018.03.008

13. Kruidhof HM, Pashalidou FG, Fatouros NE, Figueroa IA, Vet LE, Smid HM, Huigens ME. Reward value determines memory consolidation in parasitic wasps. PLoS One 2012; 7 : e39615. doi: 10.1371/journal.pone.0039615

14. Colazza S, Peri E, Salerno G, Conti E. Host Searching by Egg Parasitoids: Exploitation of Host Chemical Cues. In Consoli FL, Parra JRP, Zucchi RA, editors. Egg Parasitoids in Agroecosystems with Emphasis on Trichogramma Dordrecht, Netherlands : Springer; 2010. pp. 97–147. doi : org/10.1007/978-1-4020-9110-0_4

15. Peri E, Salerno G, Slimani T, Frati F, Conti E, Colazza S, Cusumano A. The response of an egg parasitoid to substrate-borne semiochemicals is affected by previous experience. Sci Rep. 2016; 6: 27098. doi : 10.1038/srep27098

16. Abram PK, Cusumano A, Abram K, Colazza S, Peri E. Testing the habituation assumption underlying models of parasitoid foraging behavior. PeerJ. 2017; 5: e3097. doi: 10.7717/peerj.3097

17. Schlaepfer MA, Sherman PW, Blossey B, Runge MC. Introduced species as evolutionary traps. Ecol Letters. 2005; 8: 241–246. doi: 10.1111/j.1461-0248.2005.00730.x

18. Berthon K. How do native species respond to invaders? Mechanistic and trait-based perspectives. Biol Invasions. 2015; 17: 2199–2211. doi : 10.1111/j.1461-0248.2005.00730.x

19. Cameron EK, Vila M, Cabeza M. Global meta-analysis of the impacts of terrestrial invertebrate invaders on species, communities ans ecosystems. Glob Ecol Biogeogr. 2016; 25: 596–606. doi : 10.1111/geb.12436

20. Bradley BA, Laginhas BB, Whitlock R, Allen JM, Bates AE, Bernatchez G, et al. Disentangling the abundance-impact relationship for invasive species. P Natl Acad Sci. 2019; 116: 9919–9924. doi: 10.1073/pnas.1818081116

21. Hoogendoorn M, Heimpel GE. Indirect interactions between an introduced and native ladybird beetle species mediated by a shared parasitoid. Biol Control. 2002; 25: 224–230. doi : 10.1016/S1049-9644(02)00101-9

22. Schlaepfer MA, Runge MC, Sherman PW. Ecological and evolutionary traps. Trends Ecol Evol. 2002; 17: 474–480. doi : 10.1016/S0169-5347(02)02580-6

23. Robertson BA, Rehage JS, Sih A. Ecological novelty and the emergence of evolutionary traps. Trends Ecol Evol. 2013; 28: 552–560. doi : 10.1016/j.tree.2013.04.004

24. Yeargan KV. Parasitism and predation of stink bug eggs in soybean and alfalfa fields. Environ Entomol. 1979; 8: 715–719. doi : 10.1093/ee/8.4.715

25. Koppel AL, Hebert DA, Kuhar TP, Kamminga K. Survey of stink bug (Hemiptera: Pentatomidae) egg parasitoids in wheat, soybean, and vegetable crops in southeast Virginia. Environ Entomol. 2009; 38: 375–379. doi : 10.1603/022.038.0209

26. Abram PK, Gariepy TD, Boivin G, Brodeur J. An invasive stink bug as an evolutionary trap for an indigenous egg parasitoid. Biol Invasions. 2014; 16: 1387–1395. doi : 10.1007/s10530-013-0576-y

27. Hoebeke ER, Carter ME. Halyomorpha halys (Stahl) (Heteroptera: Pentatomidae): A polyphagous plant pest from Asia newly detected in North America. Proc Entomol Soc Washington. 2003; 105: 225–237.

28. Jones AL, Jennings DE, Hooks CR, Shrewsbury PM. Sentinel eggs underestimate rates of parasitism of the exotic brown marmorated stink bug, *Halyomorpha halys*. Biol Control. 2014; 78: 61–66. doi : org/10.1016/j.biocontrol.2014.07.011

29. Haye T, Fischer S, Zhang J, Gariepy T. Can native egg parasitoids adopt the invasive brown marmorated stink bug, *Halyomorpha halys* (Heteroptera: Pentatomidae), in Europe? J Pest Sci. 2015; 88: 693–705. doi : 10.1007/s10340-015-0671-1

30. Cornelius ML, Dieckhoff C, Vinyard BT, Hoelmer KA. Parasitism and Predation on Sentinel Egg Masses of the Brown Marmorated Stink Bug (Hemiptera: Pentatomidae) in Three Vegetable Crops: Importance of Dissections for Evaluating the Impact of Native Parasitoids on an Exotic Pest. Environ Entomol. 2016; 45: 1536–1542. doi : 10.1093/ee/nvw134

31. Konopka JK, Poinapen D, Gariepy T, McNeil JN. Understanding the mismatch between behaviour and development in a novel host-parasitoid association. Sci Rep. 2018; 8: 15677. doi : 10.1038/s41598-018-33756-6

32. Gariepy TD, Bruin A, Konopka J, Scott-Dupree C, Fraser H, Bon M-C, Talamas E. A modified DNA barcode approach to define trophic interactions between native and exotic pentatomids and their parasitoids. Mol Ecol. 2019; 28: 456–470. doi : 10.1111/mec.14868

33. Costi E, Wong W, Cossentine J, Acheampong S, Maistrello L, Haye T, Talamas EJ, Abram PK. Variation in levels of acceptance, developmental success, and abortion of *Halyomorpha halys* eggs by native North American parasitoids. Biol Control. 2020; Forthcoming.

34. Tognon R, Aldrich JR, Sant’Ana J, Zalom FG. Conditioning Native *Telenomus* and *Trissolcus* (Hymenoptera: Scelionidae) egg parasitoids to recognize the exotic brown marmorated stink bug (Heteroptera: Pentatomidae: *Halyomorpha halys*). Environ Entomol. 2019; 48: 211–218. doi : 10.1093/ee/nvy186

35. Conti E, Salerno G, Bin F, Vinson SB. The role of host semiochemicals in parasitoid specificity: a case study with *Trissolcus brochymenae* and *Trissolcus simoni* on pentatomid bugs. Biol Control. 2004; 29: 435–443. doi : 10.1016/j.biocontrol.2003.08.009

36. Conti E, Salerno G, Bin F, Williams HJ, Vinson SB. Chemical Cues from *Murgantia histrionica* eliciting host location and recognition in the egg parasitoid *Trissolcus brochymenae*. J Chem Ecol. 2003; 29: 115–130. doi : 10.1023/a:1021980614512

37. Borges M, Colazza S, Ramirez-Lucas P, Chauhan KR, Moraes MCB, Richard Aldrich J. Kairomonal effect of walking traces from *Euschistus heros* (Heteroptera: Pentatomidae) on two strains of *Telenomus podisi* (Hymenoptera: Scelionidae). Physiol Entomol. 2003; 28: 349–355. doi : 10.1111/j.1365-3032.2003.00350.x

38. R Core Team. R: a language and environment for statistical computing. R Foundation for Statistical Computing; 2020. Available from : https://www.R-project.org/

39. Smid HM, Vet LEM. The complexity of learning, memory and neural processes in an evolutionary ecological context. Curr Opin Insect Sci. 2016; 15: 61–69. doi : 10.1016/j.cois.2016.03.008

40. Guerra-Grenier E, Abram PK, Brodeur J. Asymmetries affecting structure and outcome of aggressive contests between solitary egg parasitoids: the effect of natal host species. Behav Ecol. 2020; Forthcoming.

41. König K, Krimmer E, Brose S, Gantert C, Buschlüter I, König C, Klopfstein S, Wendt I, Baur H, Krogmann L, Steidle LM. Does early learning drive ecological divergence during speciation processes in parasitoid wasps? Proc R Soc B. 2015; 282: 20141850. doi : 10.1098/rspb.2014.1850

42. Vet LEM. Parasitoid foraging: the importance of variation in individual behaviour for population dynamics. In RB Floyd, editor. The Nicholson Centenary Meeting: Frontiers of population ecology. Melbourne, Australia : CSIRO Publishing; 1996. pp. 254–256.

43. Botch PS, Delfosse ES. Host-Acceptance behavior of *Trissolcus japonicus* (Hymenoptera: Scelionidae) reared on the invasive *Halyomorpha halys* (Heteroptera: Pentatomidae) and nontarget species. Environ Entomol. 2018; 47: 403–411. doi : 10.1093/ee/nvy014

44. Boyle SM, Weber DC, Hough-Goldstein J, Hoelmer KA. Parental host species affects behavior and parasitism by the pentatomid egg parasitoid, *Trissolcus japonicus* (Hymenoptera: Scelionidae). Biol Control. 2020; 149: 104324. doi : 10.1016/j.biocontrol.2020.104324

45. Heimpel G, Neuhauser C, Hoogendoorn M. Effects of parasitoid fecundity and host resistance on indirect interactions among hosts sharing a parasitoid. Ecol Lett. 2003; 6: 556–566. doi : 10.1046/j.1461-0248.2003.00466.x

46. Robertson BA, Blumstein DT. How to disarm an evolutionary trap. Conserv Sci Pract. 2019; 1: e116. doi : 10.1111/esp2.116

